# Molecular and structural basis of the heterochromatin-specific chromatin remodeling activity by *Arabidopsis* DDM1

**DOI:** 10.1101/2023.07.10.548306

**Authors:** Akihisa Osakabe, Yoshimasa Takizawa, Naoki Horikoshi, Suguru Hatazawa, Lumi Negishi, Frédéric Berger, Tetsuji Kakutani, Hitoshi Kurumizaka

**Author notes:** These two authors contributed equally to this work (Akihisa Osakabe, Yoshimasa Takizawa).

## Abstract

The chromatin remodeler DECREASE IN DNA METHYLATION 1 (DDM1) deposits the histone H2A variant H2A.W and silences transposons in *Arabidopsis thaliana*. However, the molecular mechanisms by which DDM1 specifically targets the nucleosome containing H2A.W and allows chromatin writers to access heterochromatin remained elusive. Here, we show that DDM1 promotes remodeling of the H2A.W nucleosome and requires interactions with the H2A.W-specific C-terminal tail. The cryo-EM structure of the DDM1-H2A.W nucleosome complex revealed that DDM1 binds to the N-terminal tail of H4 and the nucleosomal DNA. Comparison with the cryo-EM structure of the nucleosome containing H2A.W suggested that DDM1 increases the DNA end flexibility of nucleosomes. Based on these biochemical and structural results, we propose that the chromatin remodeling activity of DDM1 with the heterochromatin-specific H2A.W contributes to the maintenance of repressive epigenetic marks in heterochromatin by providing DNA methyltransferases with access to nucleosomal DNA.

## Introduction

Transposons are mobile DNA elements that contribute to genetic evolution in animals and plants^1–11^. The expression of transposons that results in their mobilization (referred to as transposition) can potentially disrupt gene function and threaten the integrity of the host genome. Therefore, transposons are generally silenced by heterochromatin formation with the contributions of repressive epigenetic modifications, such as cytosine methylation within DNA and the methylations of histone H3K9 and H3K27^12–14^. In plants, several DNA methyltransferases methylate DNA in CG and non-CG contexts, and DNA methylation provides a forward feedback loop to recruit H3K9 methyltransferases^15–23^, which participate in transposon silencing.

In eukaryotes, paralogs of each core histone H2A, H2B, and H3 have been identified as histone variants^24, 25^, with characteristic genome distributions, production patterns, and functions^26–33^. In *Arabidopsis*, four classes of H2A variants have been identified, and they occupy particular functional chromatin domains^34^. Among them, H2A.W evolved in land plants and is specifically localized in pericentromeric heterochromatin^34, 35^. H2A.W is distinguished from other H2A variants by its unique C-terminal tail containing the KSPKK motif, which stabilizes heterochromatin via interactions with linker DNA^34, 36–38^, and it cooperates with H3 lysine 9 dimethylation (H3K9me2) to silence transposons^39^.

Chromatin remodeling factors are proteins harboring an ATPase domain and promote the translocation of nucleosomal DNA by distorting histone-DNA interactions. Consequently, the DNA sliding and histone removal/exchange/replacement are facilitated within the nucleosome^40, 41^. In *Arabidopsis*, 41 proteins belong to the family of sucrose non-fermenting 2 (Snf2) chromatin remodeling factors^42^. Their functions are associated with various physiological pathways, including the control of flowering, flower development, and resistance to pathogens^43–52^. An Snf2-type chromatin remodeling factor named DECREASE IN DNA METHYLATION 1 (DDM1) reportedly functions in DNA methylation maintenance^53–56^ and transposon silencing^57–62^ in the *Arabidopsis* genome. DDM1 specifically binds H2A.W and mediates its deposition over transposons for silencing^35, 63^. However, the synergistic action of H2A.W and H3K9me2 accounts for the silencing of less than half of the transposons that are silenced by DDM1^39^, suggesting additional modes of action for DDM1, such as allowing DNA methyltransferases to access heterochromatin^64, 65^. Nucleosomal DNA is highly protected against DNA methylation^66, 67^, and thus the chromatin remodeling activity by DDM1 would be required for DNA methylation in the context of heterochromatin. Other than the deposition of H2A.W, in the absence of further biochemical and structural studies, the mechanisms by which DDM1 functions in transposon silencing have remained unclear.

We now report that DDM1 preferentially promotes nucleosome remodeling with the nucleosome containing H2A.W. Crosslinking mass spectrometry and biochemical analyses revealed that the H2A.W-specific C-terminal tail directly contacts DDM1 and functions in its nucleosome remodeling activity. Using cryo-electron microscopy (cryo-EM), we compared the structures of the H2A.W nucleosome and its complex with DDM1. Our analyses demonstrated that DDM1 contacts the N-terminal tail of H4, binds to the DNA within the H2A.W nucleosome, and renders the entry/exit DNA regions flexibly disordered in the H2A.W nucleosome. Based on these results, we propose that the cooperative regulation by DDM1 and H2A.W enables chromatin modifiers that deposit repressive marks to access transposons and promote their silencing in heterochromatin.

## Results

### DDM1 preferentially slides the H2A.W nucleosome

In wild type plants, H2A.W is enriched in the pericentromeric heterochromatin^34, 68^. Previous reports have shown that DDM1 changes the DNA register in nucleosomes by its nucleosome sliding activity^69^ and functions in the maintenance of DNA methylation in pericentromeric regions^65^. These findings prompted us to postulate that the H2A.W nucleosome is a preferred target of the nucleosome remodeling by DDM1. To test this hypothesis, we reconstituted nucleosomes composed of a 169 base-pair DNA fragment with the Widom 601 sequence^70^ and canonical H2A (hereafter referred to as H2A) or H2A.W, together with *Arabidopsis* histones H2B, H3.1, and H4 (Supplementary Fig. 1a-c). We used the H3 variant H3.1 for the nucleosome reconstitution because it is relatively enriched in heterochromatin compared to H3.3^35, 71, 72^. We then performed the nucleosome sliding assay using the *Bsh*1236I restriction enzyme (Fig. 1a). In this assay, the DNA substrate contains three *Bsh*1236I sites, and the nucleosome formation protects two of them (Supplementary Fig. 1a and Fig. 1). As a result, the first *Bsh*1236I site located at the linker DNA region becomes the only available cleavage site for *Bsh*1236I in the reconstituted nucleosome. However, this accessible *Bsh*1236I site could be concealed if the nucleosome sliding by DDM1 occurs (Fig. 1a). As shown in Fig. 1b, DDM1 promotes nucleosome sliding in an ATPase-dependent manner, as previously reported^69^. We found that the amounts of uncleaved DNA fragments corresponding to the nucleosome sliding were drastically higher with H2A.W nucleosomes than with H2A nucleosomes (Fig. 1b). Therefore, the H2A.W nucleosome is the preferred substrate for the nucleosome sliding activity of DDM1.

**Fig. 1:**
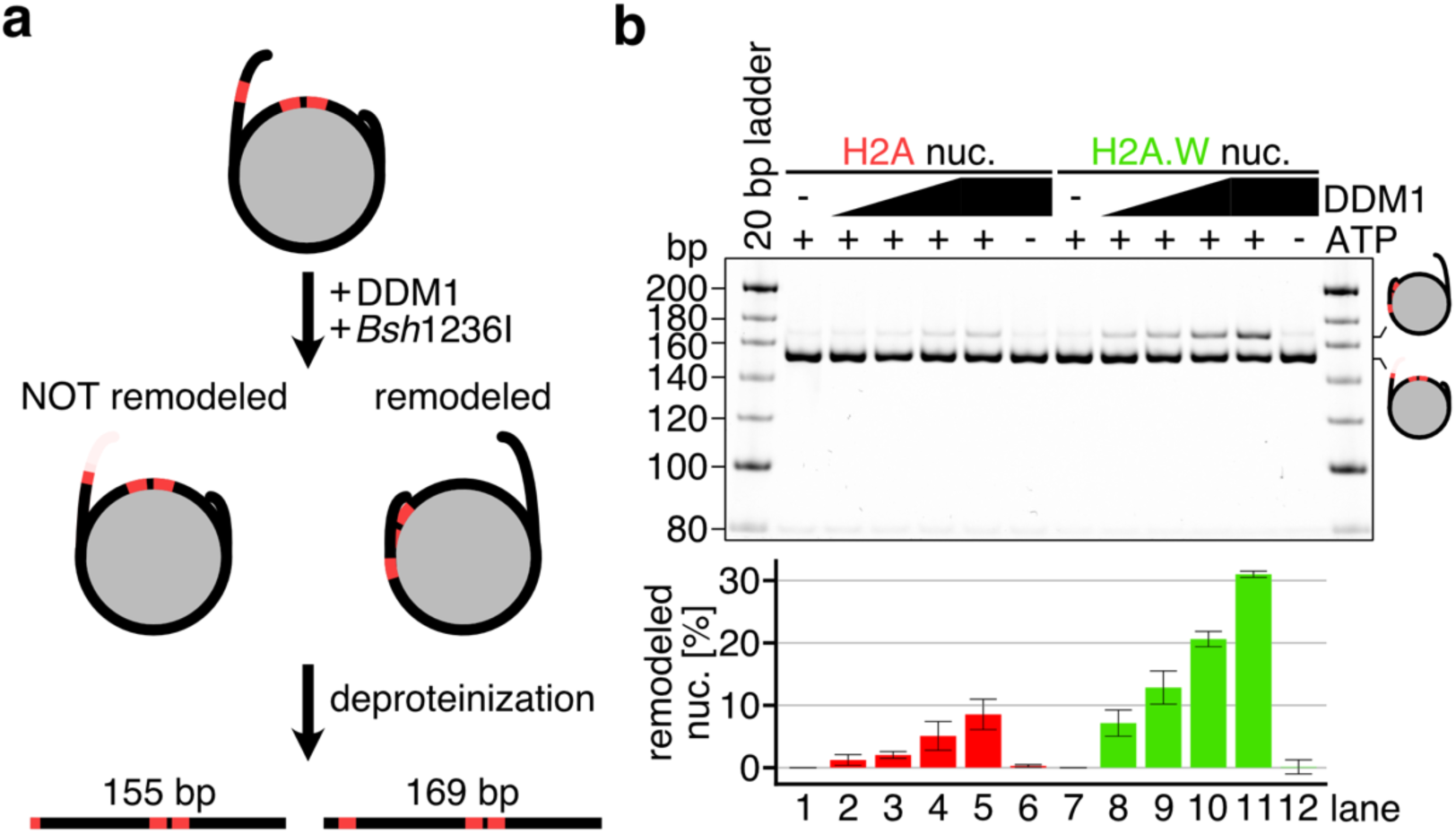
AtDDM1 preferentially slides nucleosomes containing AtH2A.W. **a,** Schematic representation of the nucleosome sliding assay using the restriction enzyme *Bsh*1236I. The sequence of the DNA fragment used in this study is provided in Supplementary Fig. 1a. **b,** Native-PAGE analyses (top panel) of DNA fragments after the nucleosome sliding assay with nucleosomes containing replicative AtH2A (red) and AtH2A.W (green). Graphical presentation (bottom panel) of the nucleosome sliding assay results. Means and error bars represent SD from three independent experiments.

### Cryo-EM structure of the DDM1-H2A.W nucleosome complex

We next determined the cryo-EM structure of the H2A.W nucleosome complexed with DDM1 (Supplementary Table 1, and Fig. 2a and b). We performed a single-particle cryo-EM workflow on the DDM1-nucleosome complex. Three-dimensional (3D) classifications identified the class of DDM1 bound to the H2A.W nucleosome, and the DDM1-H2A.W nucleosome structure was determined at 4.7 Å resolution (Supplementary Fig. 2). The central nucleosomal DNA region is located on the dyad axis of the nucleosome, and termed superhelical location (SHL0). The nucleosomal DNA locations are named every 10 base pairs from SHL0, as SHL±1, SHL±2, SHL±3, SHL±4, SHL±5, SHL±6, and SHL±7. In the DDM1-H2A.W nucleosome structure, DDM1 binds the nucleosomal DNA at SHL-2 and SHL+6 by crossing the superhelical DNA gyres (Fig. 2b-d). The nucleosomal DNA at SHL-2 is substantially distorted by DDM1 binding (Fig. 2d), in a similar manner to the nucleosomal DNA distortion induced by the yeast Snf2 nucleosome remodeler^73^ (Fig. 2d).

**Fig. 2:**
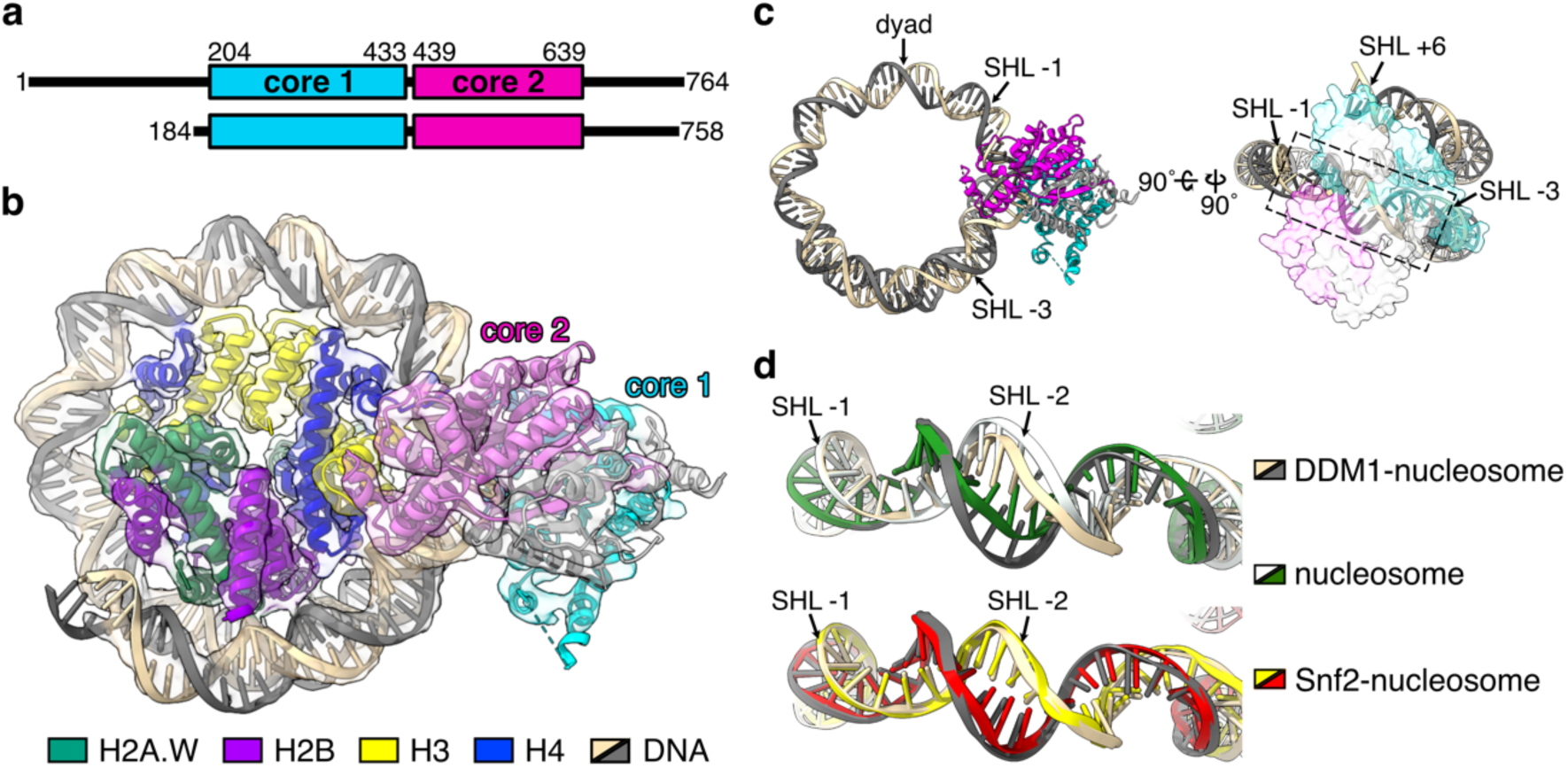
Cryo-EM structure of the AtDDM1-nucleosome complex. **a,** Schematic representation of the full-length AtDDM1 (upper) and the DDM1 fragment observed by cryo-EM (lower). The ATPase core domains 1 and 2 are colored cyan and magenta, respectively. The amino acid sequence alignment between AtDDM1 and *Saccharomyces cerevisiae* Snf2 is presented in Supplementary Fig. 3. **b,** Cryo-EM structure of the AtDDM1-nucleosome complex. The atomic structure model of the AtDDM1-nucleosome complex is fitted to the transparent cryo-EM density map. The ATPase core domains 1 and 2 of AtDDM1 are colored cyan and magenta, respectively. **c,** Structure of nucleosomal DNA bound by AtDDM1. The dashed box corresponds to the nucleosomal DNA around SHL+2, where DNA distortion was observed. **d,** Structural comparison of nucleosomal DNAs bound by AtDDM1 (light orange and grey), ScSnf2 in the absence of ADP (yellow and red, PDB ID: 5X0Y^73^), or free nucleosome (white and green).

In addition to the interaction with nucleosomal DNA, the N-terminal tail of H4 is located near the ATPase core domain of DDM1, and is captured by its acidic pocket (Fig. 3a and Supplementary Fig. 3). The binding of the H4 N-terminal tail within the acidic pocket was also reported for the yeast Snf2 protein^41, 73–75^. To test the biological significance of the H4 N-terminal tail binding to DDM1, we performed the nucleosome sliding assay with nucleosomes lacking this tail (residues 1-24). Consistent with the conserved role of the H4 N-terminal tail in the Snf2-mediated nucleosome remodeling activity^73^, the deletion of the H4 N-terminal tail drastically reduced the DDM1-mediated nucleosome sliding (Fig. 3b and Supplementary Fig. 4a-c), although weak sliding activity was still detected with the nucleosome containing H2A.W without the H4 N-terminal tail (Fig. 3b). We thus conclude that DDM1 binds and remodels nucleosomes by a mechanism similar to that described for other chromatin remodelers of the Snf2 family, but with a preference for H2A.W nucleosomes, suggesting that specific features of H2A.W enhance the chromatin remodeling activity of DDM1.

**Fig. 3:**
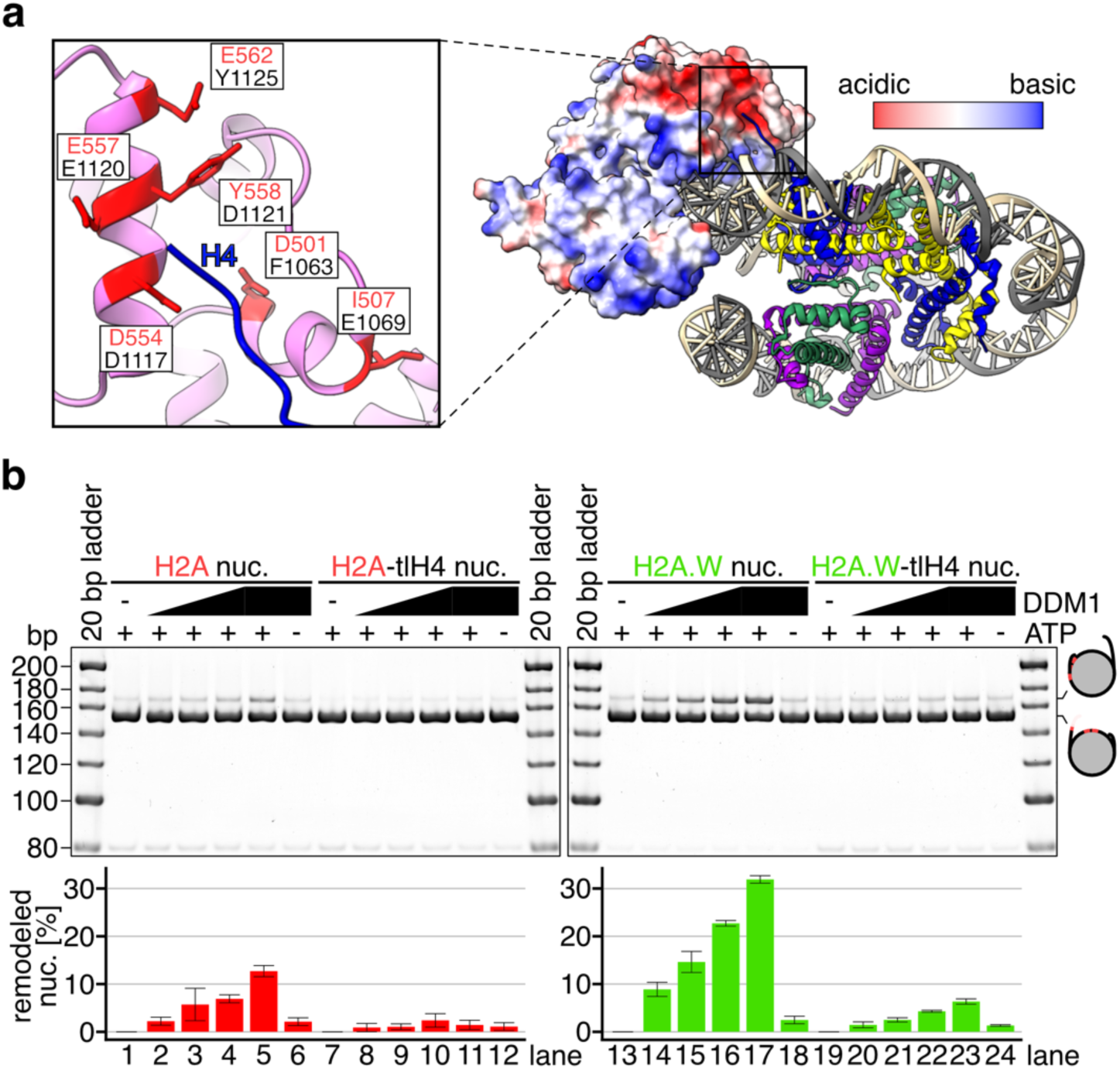
Interactions between AtDDM1 and the N-terminal tail of H4. **a,** Overall structure of the AtDDM1-nucleosome complex (right). The calculated electrostatic potential of the atomic surface of AtDDM1 molecule is presented. Close-up view of AtDDM1 and the N-terminal region of AtH4 (left). The residues of AtDDM1 surrounding the N-terminal tail of AtH4 (blue) are shown as sticks and labeled in red, where the corresponding residues of ScSnf2^73^ are also labeled in grey. The first 18 residues (aa 1-18) of AtH4 are not shown due to their flexibility. **b,** Native-PAGE analyses (top panel) of DNA fragments after the nucleosome sliding assay with nucleosomes containing AtH2A (red) and AtH2A.W (green), with or without the N-terminal tail of AtH4. Graphical presentation of the nucleosome sliding assay results (bottom panel). Means and error bars represent SD from three independent experiments.

### The C-terminal tail of H2A.W binds to DDM1 and enhances the nucleosome sliding

To identify the specific interaction between DDM1 and H2A.W, we next performed a crosslinking mass spectrometric analysis of the DDM1-nucleosome complex (Fig. 4a and Supplementary Fig. 5). We employed disuccinimidyl suberate (DSS-H12/D12), which crosslinks inter-and intra-molecular lysine residues, as the crosslinker. The profiles of DDM1-histone crosslinking are mostly conserved between the H2A.W and H2A nucleosomes (Supplementary Fig. 5). Interestingly, our crosslinking mass spectrometry analysis identified three H2A.W-specific C-terminal tails crosslinked with DDM1 residues (Fig. 4a). Two residues (K203 and K208) of DDM1 crosslinked to the C-terminal tail of H2A.W were close to regions interacting with nucleotides in the complex Snf2/nucleosomes^73, 75^. One residue (K342) was in the ATPase core region of DDM1 and faced towards the nucleosomal DNA (Fig. 4b and Supplementary Fig. 3). In our cryo-EM structure, the C-terminal tail of H2A.W may contact these residues, although the interaction between DDM1 and the H2A.W C-terminal tail was not observed, possibly due to its high flexibility (Fig. 4b). These results suggest that the C-terminal tail of H2A.W may directly bind to DDM1 and function to regulate its preferential nucleosome sliding activity.

**Fig. 4:**
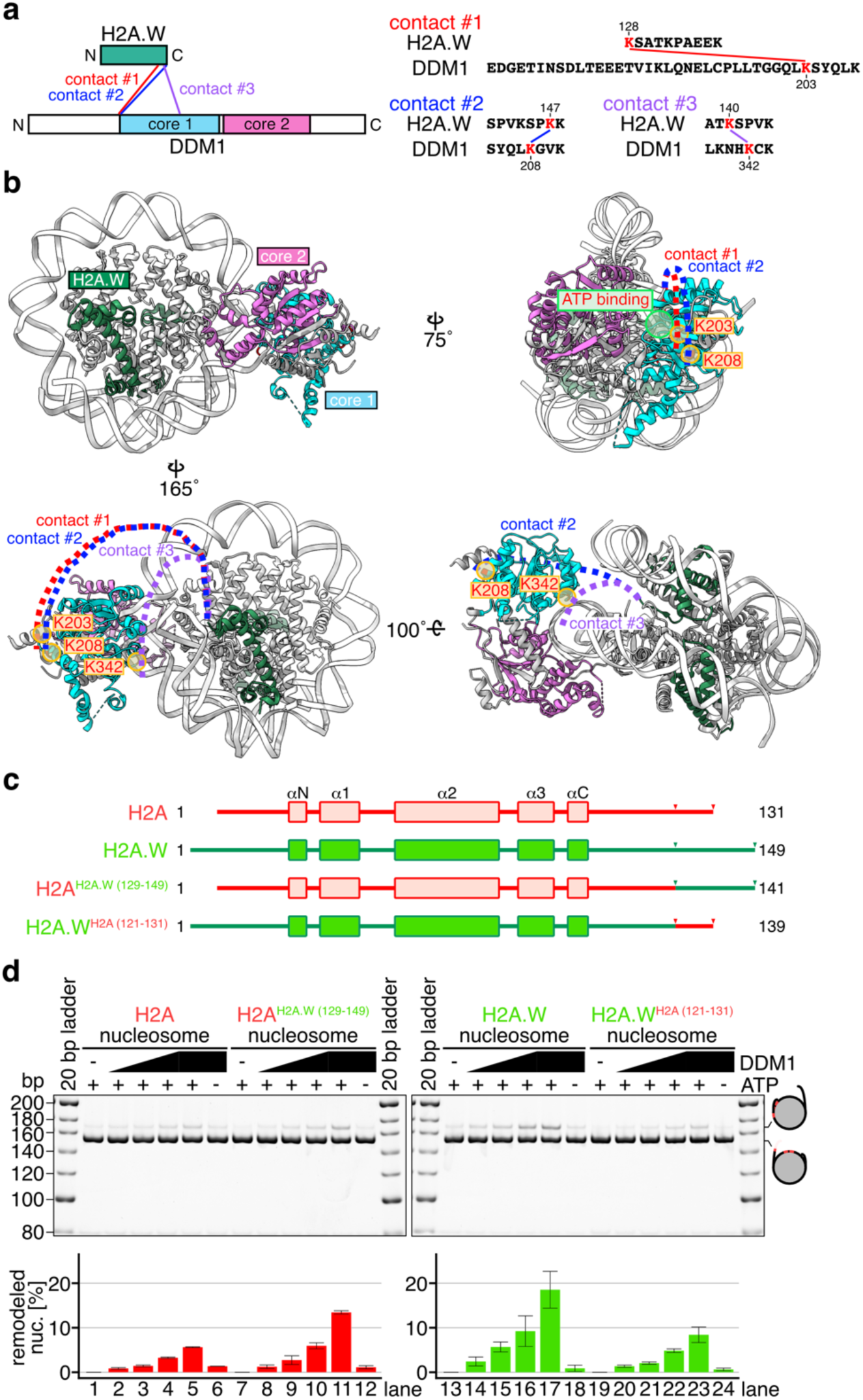
C-terminal tail of AtH2A.W contributes to enhance the nucleosome sliding activity of DDM1. **a,** Schematic representation (left) and sequence information (right) for the results obtained by crosslinking mass spectrometry of the DDM1-bound nucleosomes containing AtH2A.W. Three contacts between the H2A.W C-terminal tail and DDM1 are shown. Other crosslinks of the DDM1-bound nucleosomes containing AtH2A.W and AtH2A in the top 25% of ld-scores are shown in Supplementary Fig. 5. **b,** Graphical summary of the contacts between the H2A.W C-terminal tail and DDM1, identified by crosslinking mass spectrometry. The dashed lines indicate possible locations where the disordered H2A.W C-terminal tail contacts DDM1. The residues of DDM1 contacting the H2A.W C-terminal tail are shown in red with yellow circles. The putative ATP binding region supported by previous work^63, 75, 111^ is enclosed in a green circle. **c,** Schematic representations of the secondary structures of AtH2A, AtH2A.W, AtH2A^AtH2A.W^ ^(129–149)^, and AtH2A.W^AtH2A^ ^(121–131)^. The amino acid sequence alignment between AtH2A and AtH2A.W is shown in Supplementary Fig. 5a. **d,** Native-PAGE analyses (top panel) of DNA fragments after the nucleosome sliding assay with nucleosomes containing AtH2A and AtH2A^AtH2A.W^ ^(129–149)^ (red), AtH2A.W, and AtH2A.W^AtH2A^ ^(121–131)^ (green). Graphical presentation of the nucleosome sliding assay results (bottom panel). Means and error bars represent SD from three independent experiments.

To test this hypothesis, we prepared mutant nucleosomes containing H2A^H2A.W^ (129–149) and H2A.W^H2A^ (121–131). In H2A^H2A.W^ ^(129–149)^, the C-terminal residues 121-131 of H2A were replaced by the corresponding residues (aa 129-149) of H2A.W. Correspondingly, in H2A.W^H2A^ ^(121–131)^, the C-terminal residues 129-149 of H2A.W were switched with the corresponding residues (aa 121-131) of H2A (Fig. 4c and Supplementary Fig. 6). We then performed the nucleosome sliding assay (Fig. 4d). The DDM1-mediated nucleosome sliding was substantially enhanced with the H2A containing the C-terminal residues of H2A.W (Fig. 4d), and conversely, was drastically suppressed in the nucleosome containing H2A.W^H2A^ ^(121–131)^ (Fig. 4d). These results indicate that the H2A.W C-terminal tail plays an essential role in the nucleosome sliding mediated by DDM1. Therefore, DDM1 binds the H2A.W nucleosome through interactions with the specific H2A.W C-terminal residues, and these interactions promote nucleosome remodeling.

### DDM1 increases the flexibility of nucleosomal entry/exit DNA regions

To examine if DDM1 affects the H2A.W nucleosome structure, we determined the cryo-EM structure of the nucleosome containing H2A.W without DDM1 at 2.9 Å (Fig. 5 and Supplementary Fig. 7). This structure showed that 145 base pairs of DNA are symmetrically wrapped around the histone octamer, and 4 additional base pairs were observed as a linker (Fig. 5a-c). In contrast, in the DDM1-H2A.W nucleosome complex, only 111 base pairs of DNA are bound to the histone octamer, and the entry/exit regions from SHL-5 to SHL-7 and from SHL+6 to SHL+7 are disordered (Figs. 2 and 5). These results suggest that DDM1 binding to the H2A.W nucleosome causes the nucleosomal DNA ends to become more flexible.

**Fig. 5:**
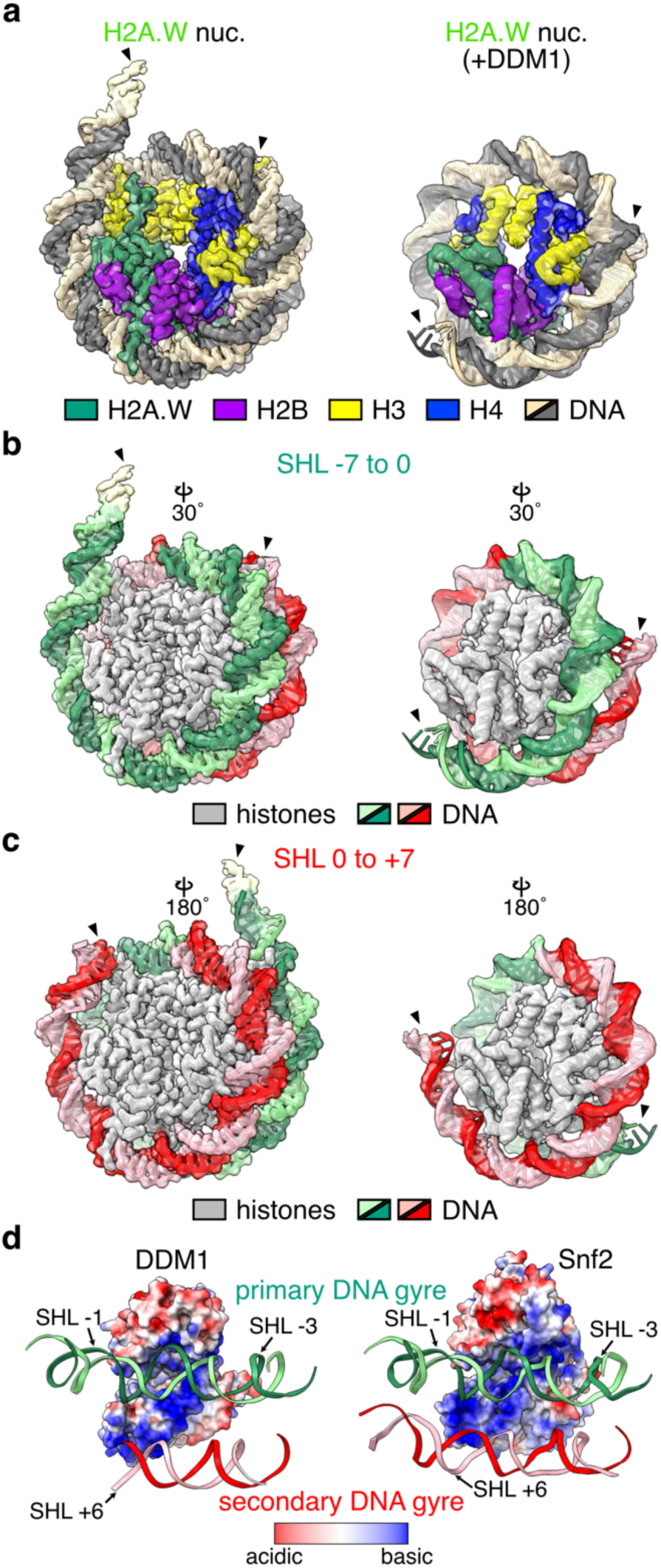
Structural comparison of DDM1-free and DDM1-bound nucleosomes. **a,** Cryo-EM structures of the nucleosome containing AtH2A.W (left), and that bound by DDM1 (right). **b and c,** Structural comparison of nucleosomal DNA at SHL -7 to 0 (**b**) and SHL 0 to +7 (**c**). Arrowheads indicate the terminal DNA detected by cryo-EM density maps. **d,** Electrostatic potentials of the atomic surfaces of DDM1 (left) and ScSnf2 (right, PDB ID: 5X0Y^73^), which face toward the primary and secondary DNA gyres.

## DISCUSSION

In the present study, we have shown that DDM1 preferentially slides H2A.W nucleosomes (Fig. 1). The H2A.W C-terminal tails and the H4 N-terminal tail play important roles in the DDM1-mediated nucleosome sliding (Figs. 3 and 4). The cryo-EM structure of the DDM1-H2A.W nucleosome complex revealed that DDM1 binds the nucleosomal DNA at the SHL-2 and SHL+6 positions (Fig. 2). In addition, a structural comparison between the DDM1-bound H2A.W nucleosome and the DDM1-free H2A.W nucleosome suggested that DDM1 increases the DNA end flexibility of nucleosomes (Fig. 5). These findings explain the mechanism by which DDM1 specifically targets the H2A.W nucleosome and promotes nucleosome remodeling to enable chromatin modifiers to access heterochromatin (Fig. 6).

**Fig. 6:**
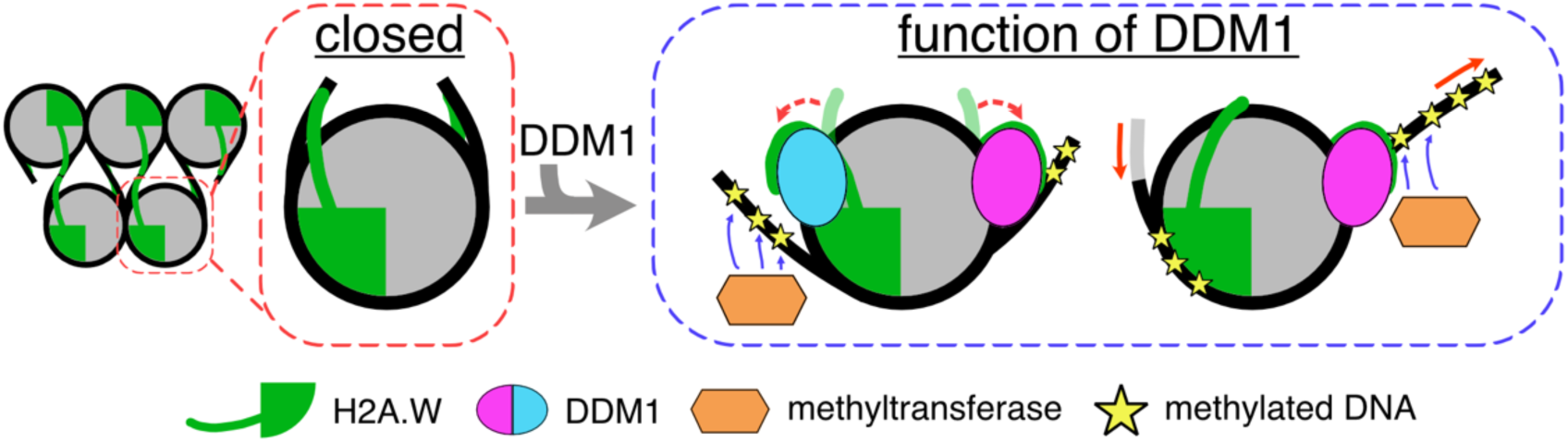
Model of DDM1 activity on the maintenance of repressive marks. In the absence of DDM1, the chromatin containing H2A.W forms a condensed structure caused by the extended C-terminal tail of H2A.W interacting with the linker DNA. In the presence of DDM1, the C-terminal tail of H2A.W might dissociate from the linker DNA by interacting with DDM1. In addition, the C-terminal tail of H2A.W contributes to enhance the remodeling activity of DDM1. These activities promote the increased accessibility of the heterochromatin to DNA methyltransferases.

Previous studies have shown that the ATPase activity of the chromatin remodeler ISWI is inhibited by its internal AutoN domain, which is enriched with basic residues and thus resembles the N-terminal tail of H4. This autoinhibition is released via competitive binding with the N-terminal tail of H4^76–79^. We noticed the sequence similarities of conserved regions in angiosperm DDM1 with the AutoN domain of ISWI and the N-terminal tail of H4 (Supplementary Fig. 4d and 4e). This implied that, like the ISWI AutoN domain, the conserved DDM1 region may also serve as a negative regulator and function via competitive binding with the H4 N-terminal tail. However, the H2A.W C-terminal tail is enriched in basic lysine residues^38, 80^, and in the present study, we found that the H2A.W C-terminal tail regulates the DDM1 activity. Our biochemical analyses showed that both the H2A.W C-terminal and H4 N-terminal tails are important for the nucleosome sliding of the H2A.W nucleosome. How these two tails function together with the DDM1 conserved region is an important issue to be addressed in the future.

In addition to our determination of the DDM1 binding mechanism to the H2A.W nucleosome, we also found that DDM1 increases the flexibility of entry/exit nucleosomal DNAs (Figs. 2 and 5). This may enhance the accessibility of DNA-binding proteins, including DNA methyltransferases. The binding of the pioneer transcription factor SOX11 to nucleosomal DNA at SHL-2 reportedly leads to a clash with the secondary DNA gyre around SHL+6, inducing the detachment of the DNAs around the entry/exit regions of the nucleosome^81^. In contrast to SOX11, DDM1 bound to SHL-2 interacts with the nucleosomal DNA around SHL+6 without steric clashes (Fig. 5d). DDM1 may directly interact with the H2A.W C-terminal tails, which are located near the DNA entry/exit regions, and the N-terminal region of H3. Therefore, DDM1 binding to the nucleosomal H2A.W C-terminal tails could perturb the histone-DNA contacts around the entry/exit regions of the nucleosome. Consistent with this idea, the cryo-EM densities are substantially ambiguous in the H2A.W C-terminal tails and the H3 N-terminal regions in the DDM1-H2A.W nucleosome complex. Further structural and biochemical analyses will be required to clarify this issue.

Our structural and biochemical data revealed the mechanism by which the chromatin remodeling factor DDM1 preferentially slides H2A.W nucleosomes. These results may provide important information for future studies of the mammalian DDM1 homolog HELLS/LSH, which mediates the deposition of a histone variant, macroH2A^82, 83^. MacroH2A is major component of heterochromatin in mammals^84^, and shares sequence similarity and functions with plant H2A.W^38^. HELLS/LSH also controls DNA methylation deposition and transposon silencing^85–87^. It is possible that HELLS/LSH preferentially remodels macroH2A nucleosomes, thus providing access to other chromatin writers. We propose that our results illustrate one of several mechanisms by which a chromatin remodeling factor cooperates with histone variants for the maintenance and epigenetic regulation of various chromatin domains in eukaryotes.

## Methods

### Preparation and purification of the nucleosome containing Arabidopsis histones

The expression and purification of the recombinant *Arabidopsis thaliana* histones were performed as described previously^37, 63, 88–90^. The DNA fragment encoding AtH2B.9 was inserted into the pET-15b vector (Novagen), in which the sequence of the thrombin recognition site (Leu-Val-Pro-Arg-Gly-Ser) was substituted with that of the TEV protease recognition site (Glu-Asn-Leu-Tyr-Phe-Gln-Gly-Ser). After purification of AtH2B.9 using nickel-nitrilotriacetic acid agarose (Ni-NTA) resin (QIAGEN), the His6-tag portion was removed by TEV protease (30 µg/mg of His6-H2B.9). AtH2B.9 was then purified on a HiPrep SP HP 16/10 cation exchange column (Cytiva), which was eluted by a linear gradient of 200-800 mM NaCl in 20 column volumes of cation exchange buffer (20 mM NaOAc, pH 5.2, 6 M urea, 5 mM 2-mercaptoethanol, 1 mM EDTA). The DNA fragment encoding truncated AtH4 (aa 25-102), AtH2A^AtH2A.W^ ^(129–149)^, or AtH2A.W^AtH2A^ ^(121–131)^ was inserted into pET-15b. Methods for the expression and purification of the truncated AtH4, AtH2A^AtH2A.W^ ^(129–149)^, or AtH2A.W^AtH2A^ ^(121–131)^ proteins were the same as described previously^37, 88–90^. The histone octamers were reconstituted and purified as described^37, 89, 90^.

The sequence of the DNA fragment used for nucleosome reconstitution is shown in Supplementary Fig. 1a. The 169 base-pair DNA fragment containing the Widom 601 sequence was purified as described previously^91^. The nucleosomes composed of the 169 base-pair DNA and histone octamers were reconstituted by the salt dialysis method, and then purified by polyacrylamide gel electrophoresis (PAGE) using a Prep Cell apparatus, as described^37, 89, 90^.

### Purification of recombinant Arabidopsis DDM1

The DNA fragment encoding AtDDM1 was inserted into the pET-15b vector (Novagen), in which the sequence of the thrombin recognition site (Leu-Val-Pro-Arg-Gly-Ser-His) was substituted with that of the human rhinovirus (HRV) 3C protease (Leu-Glu-Val-Leu-Phe-Gln-Gly-Pro), and a SUMO tag was integrated between the His6-tag and the HRV 3C protease recognition site. Expression and purification of AtDDM1 were performed by the method described previously^63^, except that GST-tagged HRV 3C protease (0.1 mg/mg of His6-SUMO-AtDDM1), instead of thrombin protease, was added to remove His6-tagged SUMO from the DDM1 portion.

### Nucleosome sliding assay

The nucleosomes (0.2 µM) were mixed with AtDDM1 (0, 0.2, 0.4, 0.8, and 1.6 µM) in a total volume of 10 µl reaction solution, containing 20 mM HEPES-NaOH (pH 7.5), 12 mM Tris-HCl (pH 7.5), 75 mM NaCl, 0.5% glycerol, 2.5 mM MgCl2, 1.2 mM DTT, 0.8 mM 2-mercaptoethanol, 0.1 mg/ml BSA and 1 mM ATP, and incubated at 30°C for 1 hour. After this incubation, the restriction enzyme *Bsh*1235I (20 units) was added to the mixtures, which were further incubated at 30°C for 30 min. After the restriction enzyme digestion, the reaction was terminated by the addition of 5 µl deproteinization solution (20 mM Tris-HCl (pH 8.0), 20 mM EDTA, 0.1% SDS, and 0.5 mg/ml Proteinase K). The resulting DNA was extracted with phenol-chloroform and then analyzed by 10% non-denaturing PAGE in 0.5 × TBE. The gel was stained with SYBR Green I solution and the DNA was visualized by an iBright Imaging System. The quantification was performed with the iBright Analysis Software (Thermo Fisher Scientific). The efficiency of the remodeled nucleosome was calculated by the ratio of the intensity of each band (169 base-pair and 155 base-pair) and normalized to the ratio obtained in the absence of AtDDM1.

### Preparation of nucleosomes and DDM1-nucleosome complexes for cryo-EM

The nucleosomes containing AtH2A.W (150 µg) were crosslinked and purified by the GraFix method^92^, using a gradient prepared with buffer A (10 mM HEPES-NaOH (pH 7.5), 1 mM DTT, and 5% (w/v) sucrose) and buffer B (10 mM HEPES-NaOH (pH 7.5), 1 mM DTT, 20% (w/v) sucrose, and 0.1% glutaraldehyde). The nucleosome containing AtH2A.W (2.39 µM) was mixed with AtDDM1 (14.34 µM) in a total volume of 300 µl reaction buffer (20 mM HEPES-NaOH (pH 7.5), 14.7 mM Tris-HCl (pH 7.5), 150 mM NaCl, 3 mM MgCl_2_, 1.2 mM DTT, 1 mM ADP, 6.2% glycerol, and 1 mM 2-mercaptoethanol), and incubated at 30°C for 30 min. Afterwards, the sample was crosslinked and fractionated by the GraFix method, using a gradient prepared with buffer A containing 150 mM NaCl and buffer B containing 150 mM NaCl. The samples were applied to the top of the gradient solution and then centrifuged at 4°C for 1 hour at 27,000 r.p.m., using an SW 41 Ti rotor (Beckman Coulter). After centrifugation, aliquots were collected from the top of the solution and analyzed by 6% (nucleosome) or 4% (AtDDM1-nucleosome complex) non-denaturing PAGE in 0.5 × TBE, followed by ethidium bromide staining. The fractions containing the nucleosome or AtDDM1-nucleosome complex were collected and then desalted on a PD-10 column (Cytiva) by elution with elution buffer (10 mM HEPES-NaOH (pH 7.5) and 2 mM TCEP). The eluted samples were concentrated using an Amicon Ultra-2 centrifuge filter unit (Merck) and stored on ice.

### Preparation of grids for cryo-EM

For the AtDDM1-nucleosome complex (0.25 mg/ml) and the nucleosome containing AtH2A.W (2.0 mg/ml), 2.5 μl portions of samples were applied onto freshly glow-discharged Quantifoil R1.2/1.3, Cu, 200-mesh grids. The grids were blotted for 8 sec at 4°C in 100% humidity, and then plunge-frozen in liquid ethane by using a Vitrobot Mark IV (Thermo Fisher Scientific).

### Cryo-EM data collection

The AtDDM1-nucleosome complex and the nucleosome containing AtH2A.W were imaged on a Krios G4 microscope (Thermo Fisher Scientific), operated at 300 kV and equipped with a BioQuantum energy filter and a K3 direct electron detector (Gatan) with a slit width of 20 eV, operated in the counting mode at a calibrated pixel size of 1.06 Å. Images of the AtDDM1-nucleosome complex and the nucleosome containing AtH2A.W were recorded at a frame rate of 150 ms for 4.5 s. A nominal defocus range of -1 to -2.5 μm was employed, and the movies were automatically acquired using the EPU software (Thermo Fisher Scientific).

### Image processing

The frames of the movies for the AtDDM1-nucleosome complex and the nucleosome containing AtH2A.W were subjected to motion correction using MOTIONCOR2, with dose weighting^93^. The contrast transfer function (CTF) was estimated using CTFFIND4^94^, and RELION4^95^ was used for the following image processing. For the AtDDM1-nucleosome complex, a total of 7,004,935 particles from dataset1 and 2,125,419 particles from dataset2 were picked by template-based auto-picking, using the 2D class averages of auto-picked particles based on a Laplacian-of-Gaussian filter as templates, followed by a few rounds of 2D classification to remove junk particles, resulting in the selection of 895,881, and 898,969 particles. The two datasets were combined, and a *de novo* initial model generated by Relion4 was low-pass filtered to 60 Å and used as the initial model for the 3D classification. The 3D class with the density map of AtDDM1 containing 411,432 particles was selected, and subjected to focused 3D classification without alignment using the AtDDM1 mask. Subsequently, 34,559 particles selected from the best classes were subjected to Bayesian polishing and CTF refinement. The final postprocessing yielded a cryo-EM map of the AtDDM1-nucleosome complex with a global resolution of 4.71 Å, with the gold standard Fourier Shell Correlation (FSC = 0.143) criteria^96^. The cryo-EM map of the AtDDM1-nucleosome complex was post-processed with the DeepEMhancer software^97^.

For the nucleosome containing AtH2A.W, 4,254,793 particles were picked by template-based auto-picking, using the 2D class averages of auto-picked particles based on a Laplacian-of-Gaussian filter as templates. After 2D classification to remove junk particles, 2,579,646 particles were selected. The crystal structure of the nucleosome^98^ (PDB ID: 3LZ0) was low-pass filtered to 60 Å and used as the initial model for the 3D classification. The selected particles were subjected to 3D classification. Subsequently, the best classes from the 3D classifications of the nucleosome containing AtH2A.W, containing 196,430 particles, were subjected to Bayesian polishing and CTF refinement. The final postprocessing yielded a cryo-EM map of the nucleosome containing AtH2A.W with a global resolution of 2.94 Å, with the gold standard Fourier Shell Correlation (FSC = 0.143) criteria^96^. The cryo-EM map of the nucleosome containing AtH2A.W was post-processed with the DeepEMhancer software^97^.

The local resolutions of the AtDDM1-nucleosome complex and the nucleosome containing AtH2A.W were calculated by RELION-4. Visualization and rendering of all cryo-EM maps were performed with UCSF ChimeraX^99^

### Model building and refinement

Model building was performed with COOT^100^, using the crystal structure of the nucleosome^98^ (PDB ID: 3LZ0) and the AtDDM1 structure generated by AlphaFold2^101^. The nucleosomal DNA was automatically fitted into the vacant volume with ISOLDE^102^. The structural models of the AtDDM1-nucleosome complex and the nucleosome containing AtH2A.W were refined by real-space refinement in Phenix^103, 104^, and validation was performed with MolProbity^105^. The data collection and statistics for the 3D reconstruction and model refinement are shown in Table 1.

### DDM1-nucleosome binding assay

The nucleosomes (0.2 µM) were mixed with AtDDM1 (0, 0.1, 0.2, 0.4, 0.8, and 1.6 µM) in a total volume of 10 µl reaction buffer, containing 20 mM HEPES-NaOH (pH 7.5), 12 mM Tris-HCl (pH 7.5), 60 mM NaCl, 0.5% glycerol, 1 mM MgCl_2_, 0.03% NP-40, 1.2 mM DTT, 0.8 mM 2-mercaptoethanol, and 1 mM ATP. The samples were incubated at 25°C for 30 min, and then analyzed by 4% non-denaturing PAGE in 0.5 × TBE (44.5 mM Tris-Borate (pH 8.3) and 1 mM EDTA). The gel was stained with SYBR Green I solution, and the DNA was visualized by an iBright Imaging System (Thermo Fisher Scientific).

### Crosslinking mass spectrometry

Nucleosomes (0.2 µM) were mixed with AtDDM1 (0.4 µM) in reaction buffer (20 mM HEPES-NaOH (pH 7.5), 60 mM NaCl, 1 mM MgCl_2_, 1.1 mM DTT, 1 mM ADP, 4.5% glycerol, 0.03% NP-40, and 0.8 mM 2-mercaptoethanol), and incubated at 25°C for 30 min. The samples were then crosslinked with 1.6 mM DSS-H12/D12 (Creative Molecules) at 25°C for 30 min, and the reaction was quenched by the addition of 50 mM Tris-HCl (pH 7.5) followed by an incubation at 25°C for 15 min. Crosslinking mass spectrometry was performed as described previously^106, 107^. The samples were dried and dissolved in an 8 M urea solution to a final protein concentration of 1.0 mg/ml. Re-dissolved samples were reduced by 2.5 mM TCEP, followed by alkylation with 5 mM iodoacetamide. For tryptic digestion, the samples were diluted with a 50 mM ammonium bicarbonate solution to a final concentration of 1 M urea, and then sequencing-grade endopeptidase Trypsin/Lys-C Mix (Promega) was added at an enzyme–substrate ratio of 1:50 wt/wt. The digested samples were applied to a Superdex 30 Increase 3.2/300 (GE Healthcare) column, using buffer containing 25% acetonitrile and 0.1% TFA. The eluted fractions (100 μl) were collected and dried completely. The residues were dissolved in 0.1% TFA and analyzed by liquid chromatography tandem mass spectrometry (LC-MS/MS), using an LTQ-Orbitrap Fusion mass spectrometer equipped with an Ultimate3000 nano-HPLC system (Thermo Fisher Scientific). The crosslinked peptides were identified using the xQuest/xProphet software^106^, and the crosslinks listed in the top 25% of ld-scores were visualized using the webserver xVis^108^.

## Data availability

The cryo-EM maps and atomic models in this study have been deposited in the Electron Microscopy Data Bank and the Protein Data Bank, under the accession codes EMD-36083 and PDB ID 8J90 for the AtDDM1-nucleosome complex and EMD-36085 and PDB ID 8J92 for the nucleosome containing AtH2A.W, respectively. The raw mass spectrometry data used in this study have been deposited to the proteomeXchange Consortium (PXD043417) via the Japan ProteOme STandard (JPOST) repository (JPST0022178)^109^.

## Acknowledgements

We thank all members of the Berger, Kakutani, and Kurumizaka laboratories, and especially Mitsuo Ogasawara (Univ. Tokyo) for technical assistance with cryo-EM data collection, and Y. Iikura, M. Dacher, and Y. Takeda (Univ. Tokyo) for their assistance. This work was supported in part by JSPS KAKENHI Grant Numbers JP21K20628, JP22H05172, and JP22H05178 [to A.O.], JP23H05475 [to H.K.], JP22K06098 [to Y.T.], and 21H04977 and 23H00365 [to T.K.], Research Support Project for Life Science and Drug Discovery (BINDS) from AMED under Grant Number JP22ama121009 [to H.K.], HFSP Grant Number RGP0025/2021 [to T.K.], JST ERATO Grant Number JPMJER1901 [to H.K.], and JST PRESTO Grant Number JPMJPR20K3 [to A.O.]. This work was also supported by the Austrian Science Fund (FWF): P32054 and P33380 [to F.B.].

## Author contributions

A.O., F.B., T.K., and H.K. conceived, designed, and supervised all the work. A.O. performed all the biochemical analyses. A.O., Y.T., and N.H. prepared the DDM1-nucleosome complex for cryo-EM. A.O. and Y.T. performed cryo-EM analyses. A.O., S.H., and L.N. performed crosslinking mass spectrometry. A.O., Y.T., F.B., T.K., and H.K. prepared all figures and wrote the manuscript. All of the authors discussed the results and commented on the manuscript.

---

**Supplementary Fig. 1:**
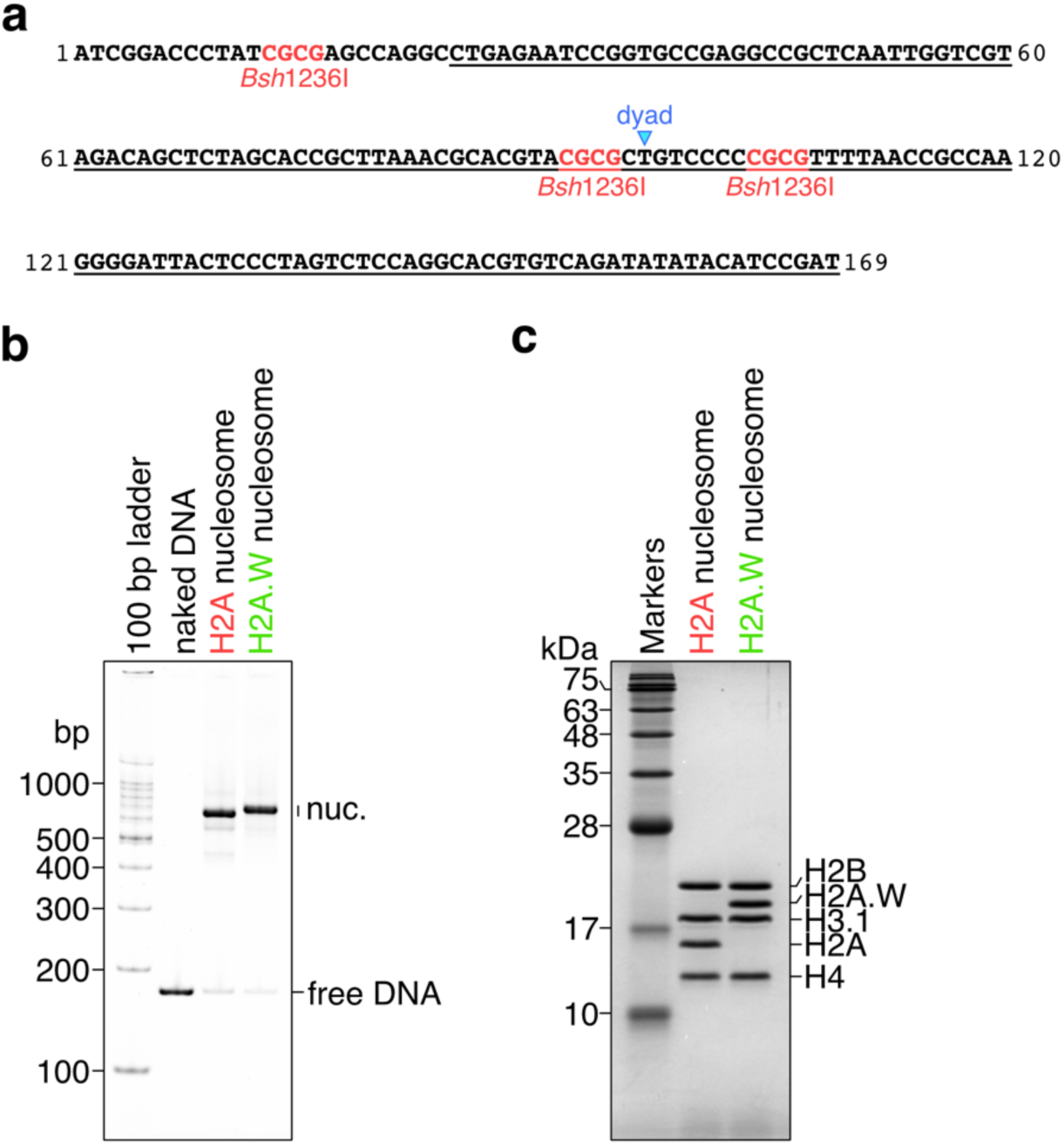
Preparation of nucleosomes containing AtH2A and AtH2A.W. **a,** Sequence of the DNA fragment used for nucleosome reconstitution. The Widom 601 positioning sequence is underlined. The blue arrowhead indicates the dyad axis. The three *Bsh*1236I recognition sites are colored red. One recognition site is 10 base-pairs away from entry/exit nucleosomal DNA and two sites are in the nucleosome positioning sequence around the dyad axis. **b and c,** Native-PAGE (**b**) and SDS-PAGE (**c**) analyses of purified nucleosomes containing AtH2A and AtH2A.W.

**Supplementary Fig. 2:**
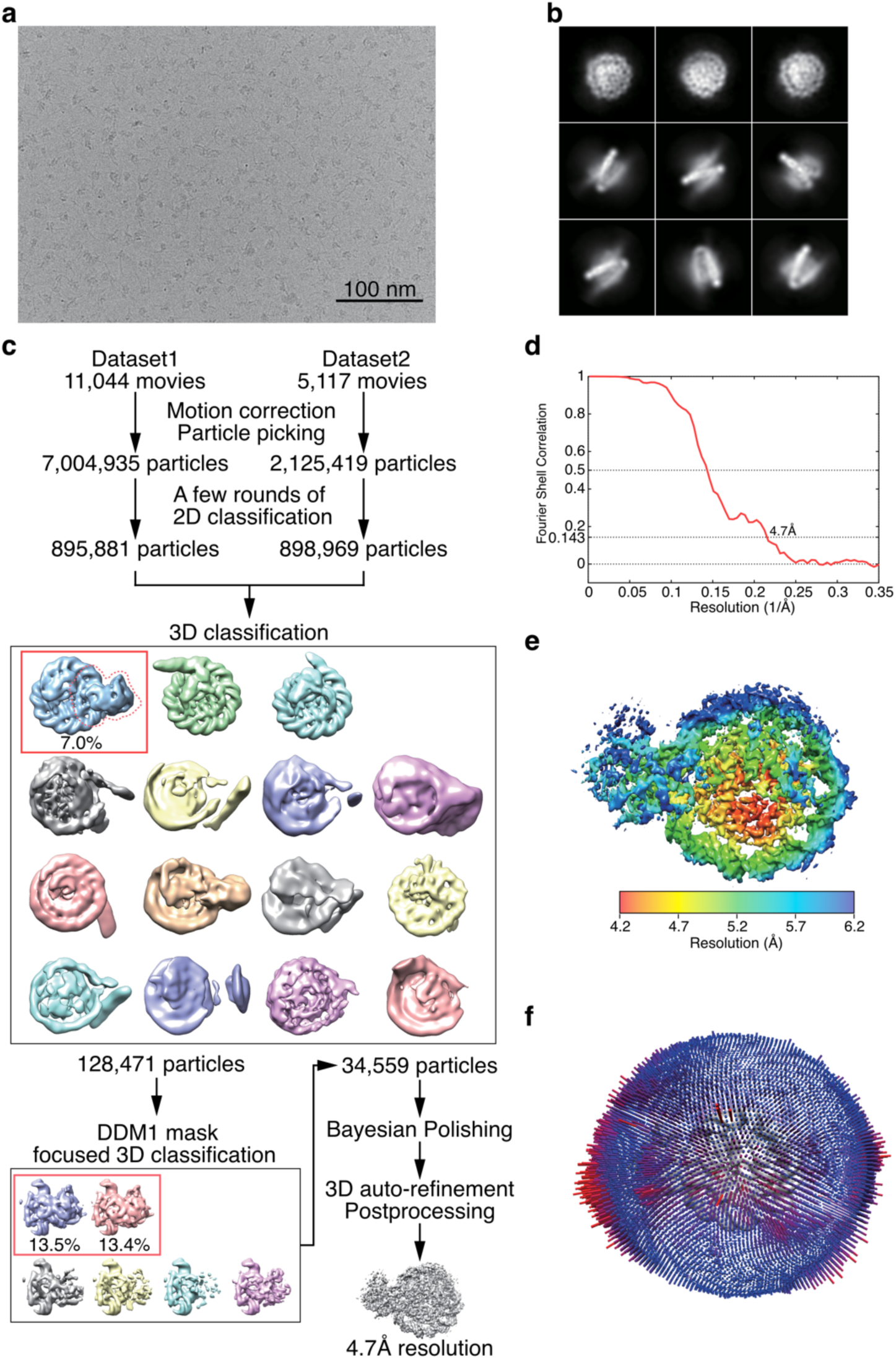
Cryo-EM data collection and image processing of AtDDM1-nucleosome complex. **a,** Representative cryo-EM micrograph. Scale bar: 100 nm. **b,** Representative 2D class averages. Box size is 21.2 nm. **c,** Flow chart of the cryo-EM image processing. **d,** The “gold standard” Fourier Shell Correlation curve calculated between 3D reconstructions from two halves of data sets. **e,** Local resolution map. **f,** Euler angle distribution of the 3D reconstruction.

**Supplementary Fig. 3:**
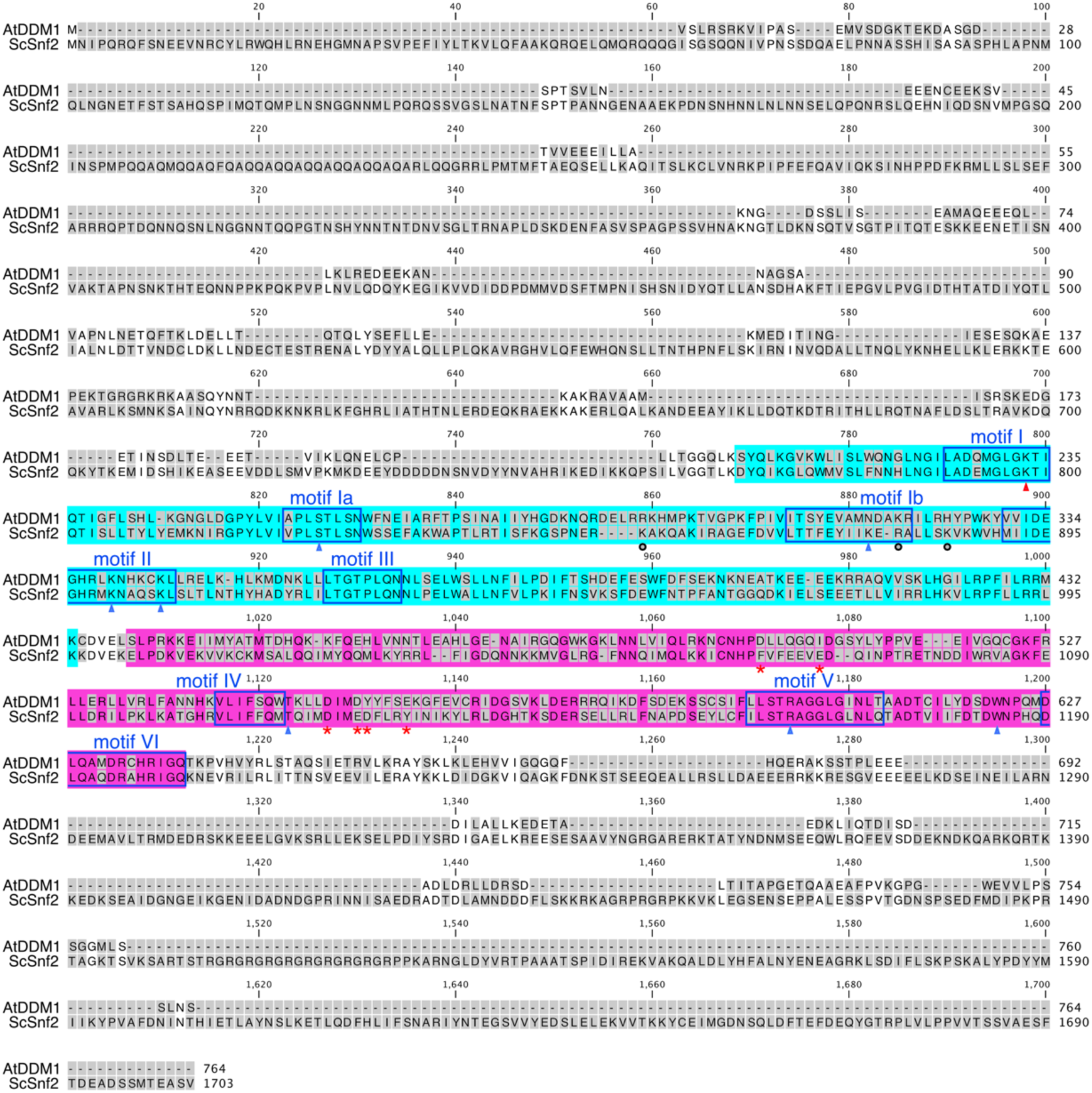
The alignment of amino acid sequences between AtDDM1 and ScSnf2. The amino acid residues that differ between AtDDM1 and ScSnf2 are shown with a grey background. The amino acid residues corresponding motifs I-VI identified in *Myceliophthora thermophila* Snf2^110^ are shown in blue boxes. Cyan and magenta boxes correspond to ATPase core domains 1 and 2, respectively. The red arrowhead indicates the residue involved in nucleotide binding, discussed in previous studies^63, 75, 111^. Blue arrowheads and black circles indicate residues involved in primary and secondary DNA binding, respectively, as discussed previously^73^. Red asterisks indicate residues that may be involved in binding to the N-terminal tail of H4, as shown in Fig. 3a.

**Supplementary Fig. 4:**
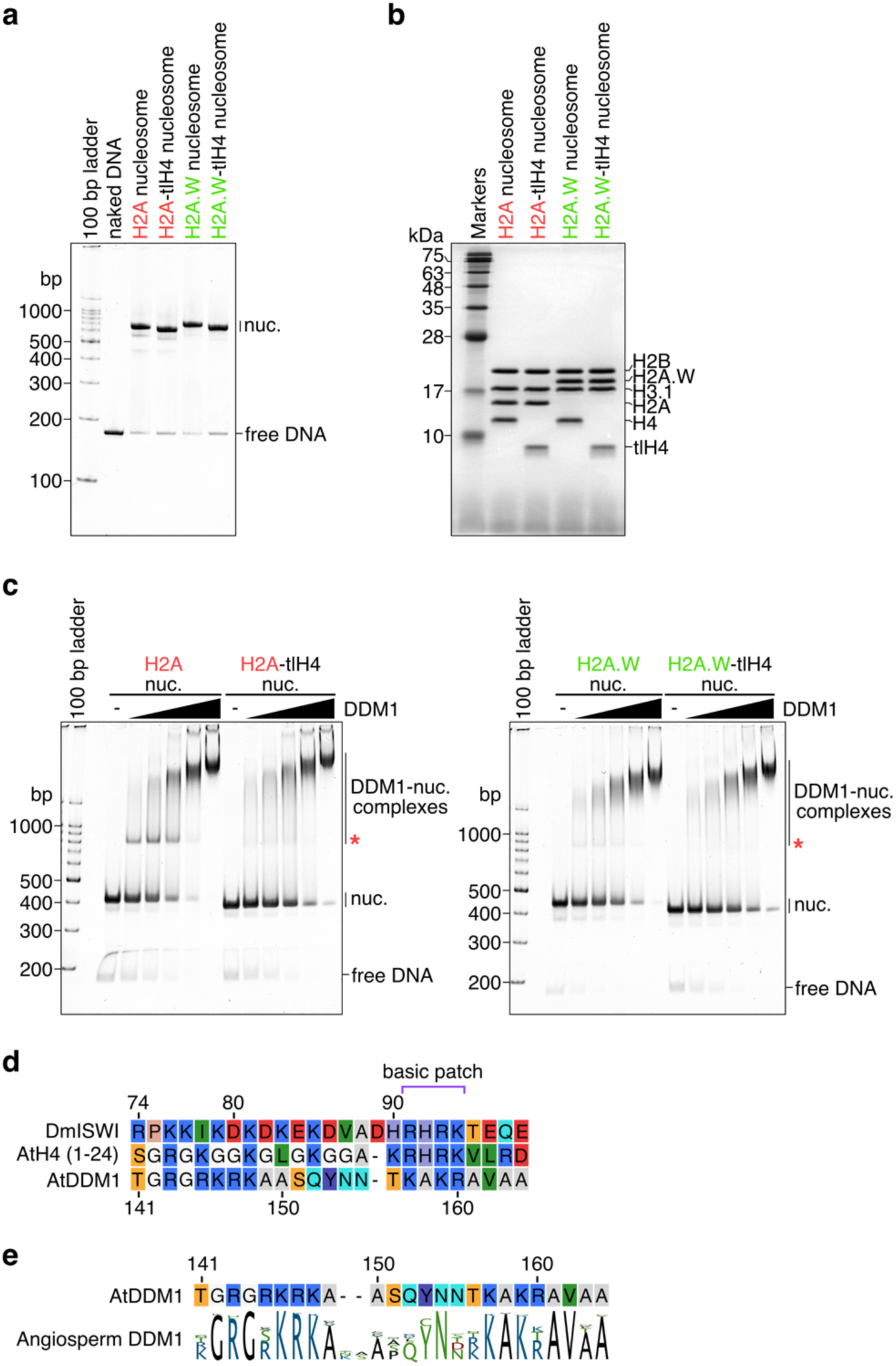
Preparation of nucleosomes lacking the N-terminal tail of AtH4. **a and b,** Native-PAGE (**a**) and SDS-PAGE (**b**) analyses of purified nucleosomes containing AtH2A and AtH2A.W, with or without the N-terminal tail of AtH4. **c,** Native-PAGE analyses of the electrophoresis mobility shift assay for the binding between AtDDM1 and nucleosomes. Specific bands corresponding to DDM1-nucleosome complexes (indicated by red asterisks) disappear in the nucleosome lacking the N-terminal tail of AtH4. **d,** Amino acid sequence alignment of the *Drosophila* ISWI AutoN region, the N-terminal tail of AtH4, and AtDDM1. The basic patch defined previously^76, 79^ is shown with a purple bracket. **e,** Sequence logo of alignments of DDM1s from flowering plants (angiosperms)^63^ with the sequence from AtDDM1.

**Supplementary Fig. 5:**
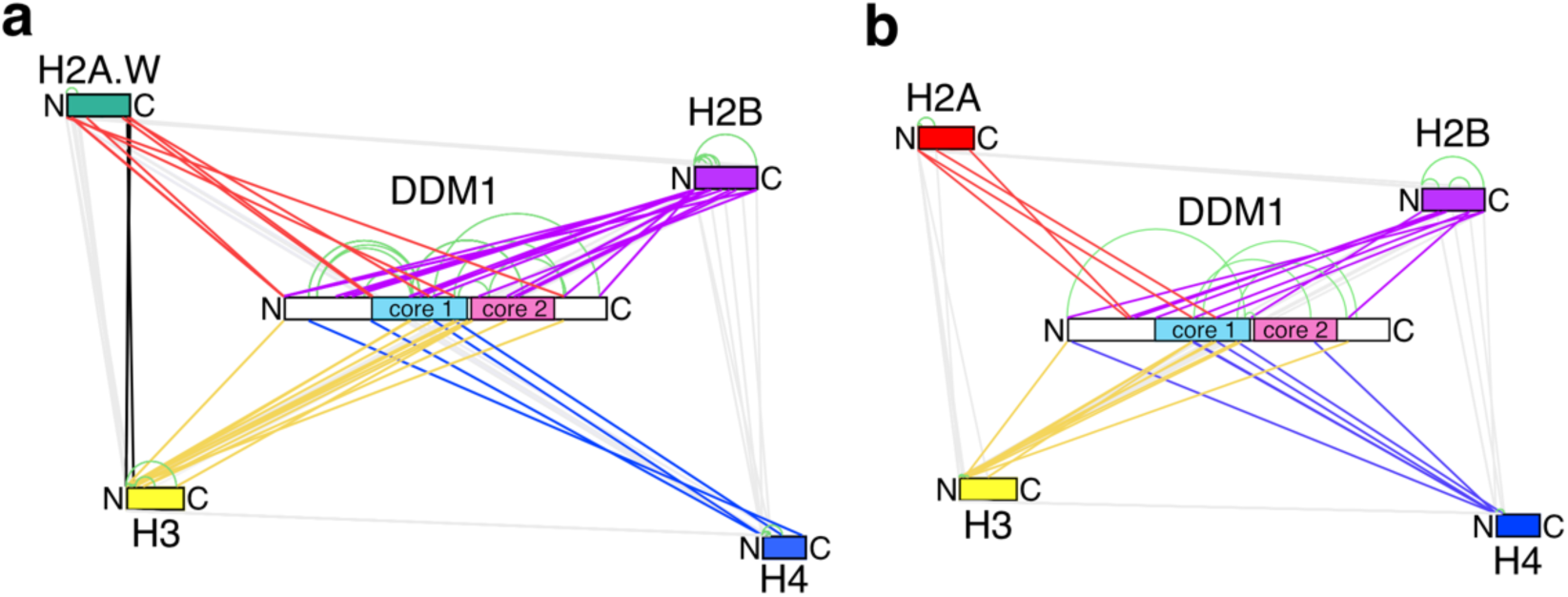
Crosslinking mass spectrometry of DDM1-bound nucleosomes. **a and b,** Schematic representation of the results obtained by crosslinking mass spectrometry of the DDM1-bound nucleosomes containing AtH2A.W (**a**) and AtH2A (**b**). Grey (between histones), black (between AtH2A.W and AtH3), red (between AtH2A or AtH2A.W and AtDDM1), purple (between AtH2B and AtDDM1), yellow (between AtH3 and AtDDM1), and blue (between AtH4 and AtDDM1) lines correspond to inter-protein crosslinks, and green lines correspond to intra-protein crosslinks.

**Supplementary Fig. 6:**
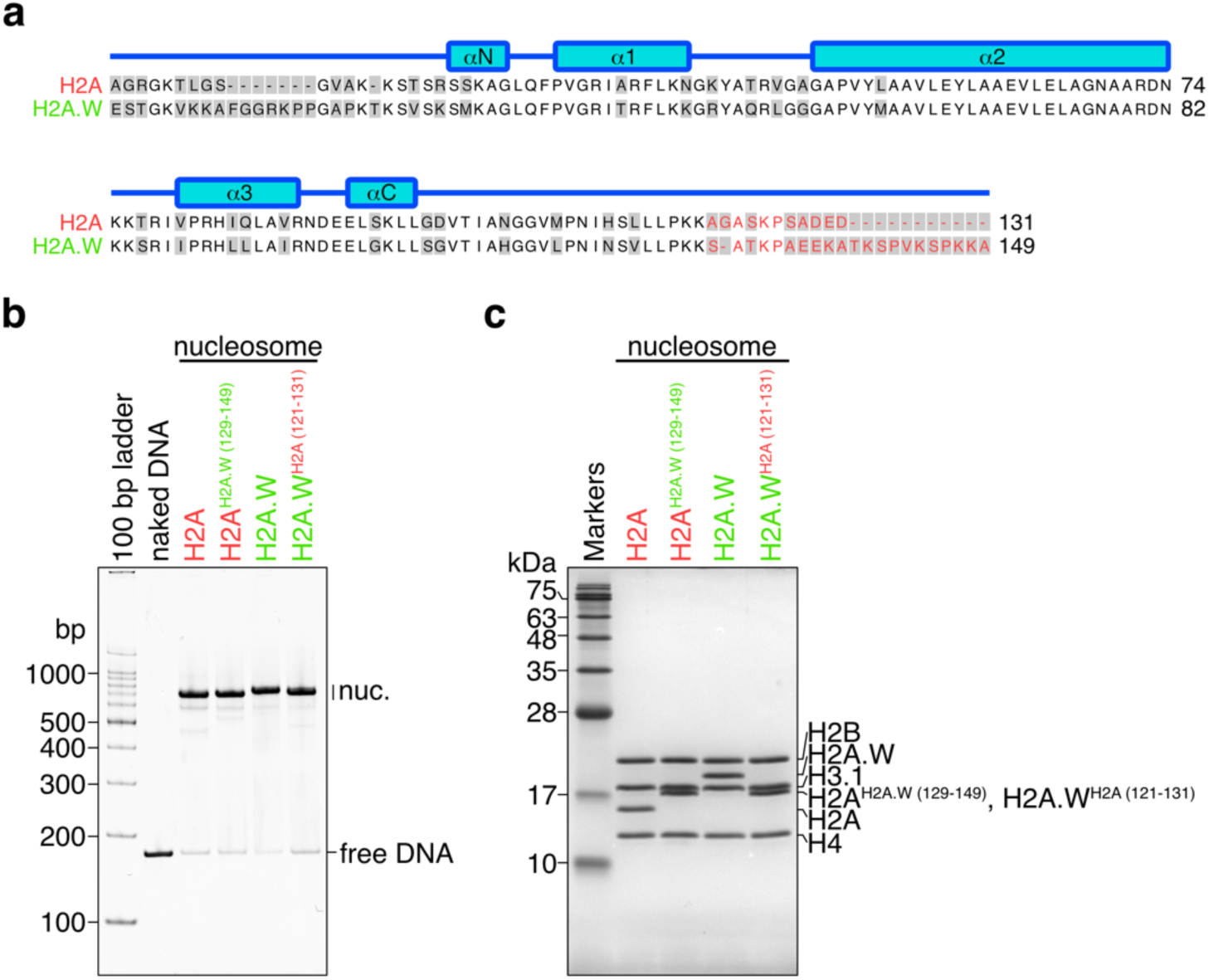
Preparation of nucleosomes containing chimeric H2A variants. **a,** Amino acid sequence alignment between AtH2A and AtH2A.W. The amino acid residues that differ between AtH2A and AtH2A.W are shown with a grey background. The red-colored amino acid residues were swapped to create the AtH2A^AtH2A.W^ ^(129–149)^ and AtH2A.W^AtH2A^ ^(121–131)^ chimeric H2A variants. **b and c,** Native-PAGE (**b**) and SDS-PAGE (**c**) analyses of the nucleosomes containing AtH2A, AtH2A^AtH2A.W^ ^(129–149)^, AtH2A.W, and AtH2A.W^AtH2A^ ^(121–131)^.

**Supplementary Fig. 7:**
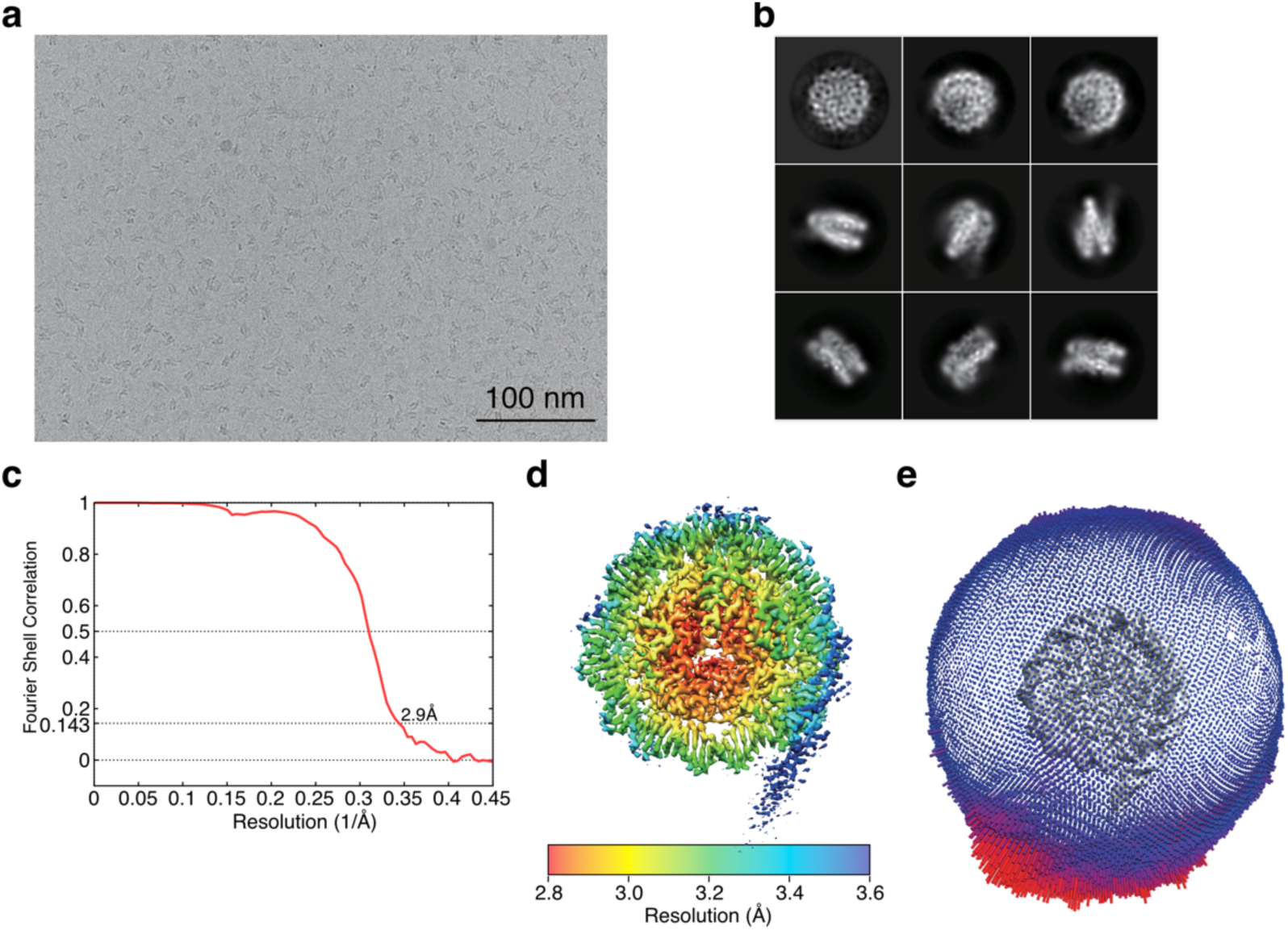
Cryo-EM data collection and image processing of nucleosomes containing AtH2A.W. **a,** Representative cryo-EM micrograph of the AtH2A.W nucleosome. Scale bar is 100 nm. **b,** Representative 2D class averages of the AtH2A.W nucleosome. Box size is 21.2 nm. **c**, The “gold standard” Fourier Shell Correlation curve calculated between the 3D reconstructions from two halves of the data set for the AtH2A.W nucleosome. **d**, Local resolution maps of the AtH2A.W nucleosome. **e**, Euler angle distributions of the AtH2A.W nucleosome.

